# Examining the Reproductive Success of Bull Kelp (*Nereocystis luetkeana*) in Climate Change Conditions

**DOI:** 10.1101/2022.11.01.514766

**Authors:** Angela R. Korabik, Tallulah Winquist, Edwin D. Grosholz, Jordan A. Hollarsmith

## Abstract

Climate change is affecting marine ecosystems in many ways including rising temperatures and ocean acidification. From 2014-2016, an extensive marine heatwave extended along the west coast of North America and had devastating effects on numerous species during this period, including bull kelp (*Nereocystis luetkeana*). Bull kelp is an important foundation species in coastal ecosystems that can be affected by marine heat waves and ocean acidification, however these impacts have not been investigated on sensitive early life stages. To determine the effects of changing temperatures and carbonate levels on Northern California’s bull kelp populations, we collected sporophylls from mature bull kelp individuals in Point Arena, CA. At the Bodega Marine Laboratory, we released spores from field-collected bull kelp, and cultured microscopic gametophytes in a common garden experiment with a fully factorial design crossing modern conditions (11.63±0.54°C and pH 7.93±0.26) with observed extreme climate conditions (15.56±0.83°C and 7.64±0.32pH). Our results found that both increased temperature and decreased pH influenced growth and egg production of bull kelp microscopic stages. Increased temperature generally resulted in decreased gametophyte survival and offspring production. In contrast, decreased pH had less of an effect, but generally resulted in increased gametophyte survival and offspring production. Additionally, we found that increased temperature significantly impacted reproductive timing by causing female gametophytes to produce offspring earlier than under ambient temperature conditions. Our findings inform better predictions of the impacts of climate change on coastal ecosystems as well as provide key insight into environmental dynamics regulating the bull kelp lifecycle.

## Introduction

Globally, marine systems are under pervasive threats from climate change. Chief among these threats are marine heat waves and ocean acidification (OA) (Cooley et al. 2022). Changing temperature and OA have negative impacts on the critical structure-forming foundational species of the world’s oceans, namely kelps and corals, especially in terms of reduced reproduction (Straub et al. 2019, Smith et al. 2022) and juvenile mortality (Kroeker et al. 2013, Harvey et al. 2013, Przeslawksi et al. 2014). In the ocean, early life stages are already subject to high mortality rates due to a number of environmental bottlenecks, and increased temperature and decreased pH can further increase juvenile mortality through reduced recruitment and growth of microscopic stage canopy kelps (Gaitan-Espitia et al. 2014, Shukla and Edwards 2017, Hollarsmith et al. 2020, Lind and Konar 2017), reduced calcification and increased disease in juvenile invertebrates (Kroeker et al. 2013, Ban et al. 2013, Small et al. 2016, Miner et al. 2018), and altered larval fish behavior (Munday 2010, Ferrari 2011).

Kelp forests are critical to temperate, nearshore subtidal and intertidal marine systems worldwide, and sustain numerous economically important recreational and commercial fisheries (Bennett et al. 2016, Blamey and Bolton 2018, Carr and Reed 2016). In addition, kelp forests provide numerous ecosystem functions and services such as shelter of structural habitat and food sources to surrounding ecosystems, buffering coastlines from wave energy, ameliorating the effects of ocean acidification, reduction of current speeds and larval delivery to the shore, and modification of seawater chemistry (Hamilton et al. 2022, Malone et al. 2022).

Globally, the effects of marine heatwaves are already having extreme effects on kelp forests (Arafeh-Dalmau et al. 2019, Straub et al. 2019, Filbee-Dexter et al. 2020). From 2014-2017, Northern California lost 90% of its bull kelp (*Nereocystis luetkeana*) kelp canopy cover over an area of roughly 350 km (Rogers-Bennett and Catton 2019). This loss of kelp forest cover has been attributed to a dramatic increase in purple urchin (*Strongylocentrotus purpuratus*) density due to loss of keystone predators, coupled with a pervasive system of marine heatwaves (McPhearson et al. 2021). The results of such widespread canopy loss were drastic changes in community structure and composition (Beas-Luna et al. 2020) and the collapse of the several fisheries in the area, such as that of the red sea urchin (*Mesocentrotus franciscanus*) (Rogers-Bennett and Okamoto 2020) and the closure of the world’s largest recreational abalone fishery (*Haliotis rufescens*) (Rogers-Bennett and Catton 2019, Reid et al. 2016).

Numerous studies in recent years have documented the effects of increased temperature on bull kelp canopies (Rogers-Bennett and Catton 2019, Hamilton et al. 2020, Helen Berry’s work in PS?), and have found that decreases in adult bull kelp canopy abundance have been related to local and large scale processes associated with warm water (Schiel et al. 2004, Pfister et al. 2017). Bull kelp exposure to warm temperatures also reduces adult blade morphological plasticity to changes in hydrodynamic flow regimes (Suprataya et al. 2020), but the physiological impacts of warm waters on bull kelp need to be further studied. Studies of bull kelp microscopic developmental stages in British Columbia and Alaska have found that increased temperatures have resulted in reductions in settlement and reduced germination and growth (Schiltroth 2021, Lind and Konar 2017, Muth et al. 2019), but the impact of rising temperatures on microscopic bull kelp stages in the southern portion of their range in northern California remains unclear. California bull kelp populations represent the range extreme of bull kelp, existing in low-latitude areas that are the most exposed to ENSO warm water events compared to more northern populations. As a result, California bull kelp populations could either be more warm-water adapted than the higher latitude populations previously studied, or they could be existing much closer to the thermal maxima and therefore very vulnerable. As the bulk of kelp die offs during the 2014 to 2017 marine heatwave occurred near the lower-latitude portion of kelp species’ ranges (Arafeh-Dalmau et al. 2019, Beas-Luna et al. 2020, Cavanaugh et al. 2019, Finger et al. 2021, Rogers-Bennett and Catton 2019), it is necessary to further study how future marine heatwaves may affect the ability of these foundation species to remain in their lower latitude ranges.

In addition to the increasing threat of marine heat waves, coastal temperate ecosystems are also subject to stress from ocean acidification (OA), which, on average, has already caused a global lowering of surface water pH by 0.11 pH units (Feely et al., 2004, Feely et al., 2009, Gattuso et al., 2015a). Variability of pH levels in nearshore systems is normal to a degree as seasonal oceanographic shifts like upwelling bring deep offshore waters to the surface and expose nearshore ecosystems to reduced pH levels. This exposure varies with local bathymetry and coastal topography, which often changes the intensity of upwelling events along the coast (Feely et al. 2008). While pH variation in the California Current System generally stays between 7.720 and 8.413 (Feely et al. 2018), climate change projections predict an increasing frequency and duration of low-pH extremes (Bakun et al. 2015, García-Reyes et al. 2015), which may result in an average decrease of up to 0.4 pH units (Feely et al. 2008). Low pH may impact physiological functions among a variety of organisms. Studies have shown that OA will more proportionately impact organisms that form calcium carbonate skeletons (Kroeker et al. 2013), but it is critical that we also understand how the compounding stress of these combined threats will impact our critical temperate nearshore systems.

Kelps are very efficient at processing multiple carbon species in the water column. Kelps utilize CO_2_ via photosynthesis, and also have the ability to concentrate calcium carbonate into their tissues. There is some evidence to suggest that the excess of carbon predicted for future ocean conditions may increase kelp growth in climate change conditions. For example, increased *p*CO_2_ has been shown to have beneficial impacts on mature bull kelp net apparent productivity (Thom 1996) and growth (Swanson and Fox 2007). At the microscopic stage, however, the effects of *p*CO2 and pH on kelp can be variable, ranging from having no effect (Fernández et al. 2015, Hollarsmith et al 2020) to positive effects on growth and photosynthesis.

Understanding how different life stages respond to environmental stress is critical when trying to predict population resilience to disturbance events. Laminariales, or the large canopy forming kelps, have a multistage process of development that presents numerous areas for the imposition of bottlenecks from climate stress. However, to our knowledge, no studies have yet investigated the role that pH may play in juvenile bull kelp development, nor the combined threats of increased temperature and ocean acidification on any bull kelp life stage.

In this study, we ask how increased temperatures and decreasing pH will impact: 1) bull kelp gametophyte development, 2) egg and sporeling production, and 3) juvenile growth. Based on the observed negative effects of the 2014-17 marine heatwave on bull kelp adults, we hypothesized that increased temperature will generally result in decreased growth, survival and reproduction. In contrast, we hypothesized that decreased pH has less of an effect than temperature on growth and egg production, but generally resulted in increased growth, survival and reproduction.

## Materials and Methods

### Bull Kelp Life Cycle

In California, one of the most dominant canopy forming kelp species is bull kelp (*Nereocystis luetkeana*). The range of bull kelp extends from the eastern Aleutian Islands, Alaska in the north to Point Conception, CA in the south. Within its California range, it is considered to be the dominant canopy-forming kelp species in Northern California, between San Francisco and the California-Oregon border. Bull kelp is an annual species, and is thought to be a more opportunistic, resilient colonizer, especially in areas too turbulent for the persistence of giant kelp (*Macrocystis pyrifera*) (Foster and Schiel 1985, Graham 1997, Graham et al. 2007).

Bull kelp have a heteromorphic life cycle consisting of a large diploid sporophyte, and a microscopic haploid gametophyte. Adult sporophytes develop sporophyll patches on their blades at the ocean surface, and at maturity, begin to release spores. The released zoospores then settle on hard substrate at the benthos, where they grow into microscopic male and female gametophytes. The female gametophytes begin to develop eggs, and then release a pheromone to trigger sperm release from nearby males. Once the sperm fertilizes the egg, a new sporophyte begins to develop (Reed 1990).

### Collection

Sporophylls from approximately 10 individuals were collected at the surface by boat from a single kelp bed in Point Arena, California (38.916271°N, 123.725644°W) in October 2017. Sporophylls were cleaned in iodine and freshwater, layered in a cooler with wet paper towels separating individual sporophylls, and transported to the Bodega Marine Laboratory (BML, 38.318164°N, 123.072019°W) for sporulation. Spore densities were determined using a hemocytometer (model number CTL-HEMM-GLDR, LW Scientific, Lawrenceville, U.S.A.), and were introduced into the experimental petri dishes at densities of approximately 8 spores/mm^2^.

### Ex-situ culturing experiment

We conducted a fully factorial common garden experiment that consisted of four treatments representing ambient and high temperature and ambient and low pH, with ten replicates per treatment, for a total of forty experimental petri dishes (Fisher Brand 100mm x15mm). Temperature was maintained at 15.6±0.8°C and 11.6±0.5°C using incubators at BML, and pH was maintained at 7.93±0.26 pH and 7.64±0.32 pH using chemical additions of equal parts 1M HCl and 1M NaHCO3 (NaHCO3 + HCl → NaCl + H2CO3) (Riebesell et al., 2011).

Temperatures were chosen to represent ambient sea surface temperatures for our ambient temperature treatment, whereas our high temperature treatment represented the 4°C increase in SST observed during the 2014-17 marine heatwave (Gentemann et al. 2017). Ambient and low pH were chosen to represent the pH of incoming seawater at BML and pH during an extreme upwelling event (Feely et al., 2008), respectively. Light was set at 14:10 photoperiod and 30-45 umol m^-2^ s^-1^. The pH of incoming, manipulated, and outgoing seawater was measured to 0.01 pH units immediately after collection using a spectrophotometer. Total alkalinity (T_alk_, μmol kg^−1^) was measured using potentiometric acid titration. Water was changed in all experimental dishes every 2 to 3 days for the duration of the 27 day experiment. We added standard 20 mL L^-1^ Provasoli nutrient mix to all treatment water (Provasoli 1968).

### Photo Analysis

Beginning one week after spore inoculation, petri dishes were photographed weekly with a Micropublisher 5.0 RTV digital camera (QImaging, Surrey, Canada) mounted on an inverted microscope at 40x magnification, resulting in 4 weeks of photos documenting population growth and reproduction. Within each dish, 3 points were randomly selected to be photographed, with different points being photographed each week. Each photo encompassed 1.08mm^2^ of the petri dish (15,708mm^2^ bottom surface area).

After the growth experiment was completed, each photo was analyzed using ImageJ (Rasband 2019). Week 1 and 2 photos did not contain any gametophytes large enough to identify by sex, so only Weeks 3 and 4 were used for analysis. Count data was obtained from each photo for female gametophytes, male gametophytes, eggs, and juvenile sporelings (Figure 1). Females were also categorized as “productive” (having produced at least one egg or juvenile) or “non-productive” (have no eggs or juveniles). The proportion of productive females in each photo was calculated by dividing the number of productive females by the total number of females counted.

**Figure 1:**
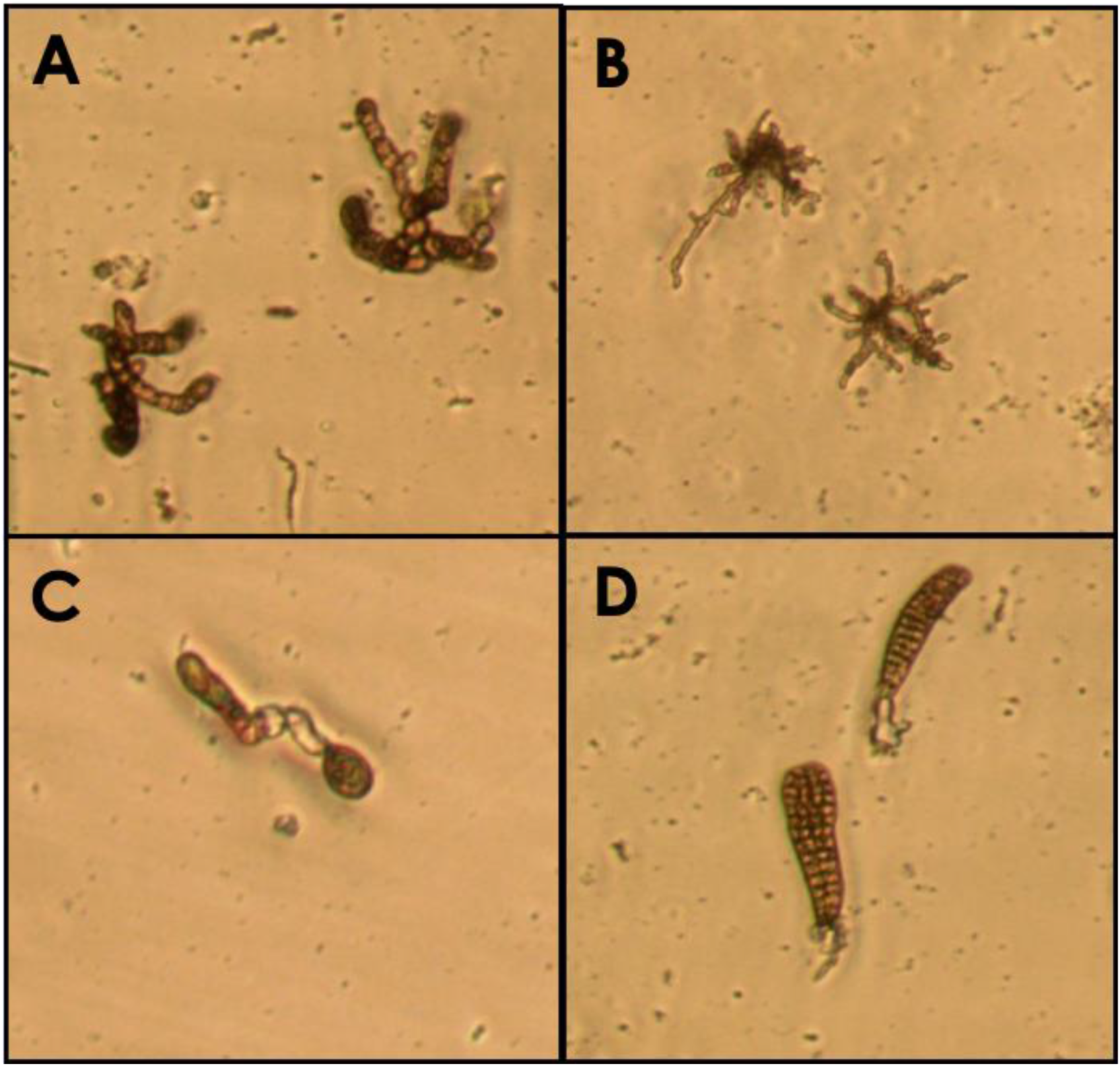
The microscopic stages of bull kelp: A) Female gametophyte (image area =0.065 mm^2^); B) Male gametophyte (image area = 0.065 mm^2^); C) Female gametophyte producing an egg (image area = 0.077 mm^2^); D) Female gametophytes with juvenile sporelings (image area = 0.065 mm^2^).

Count data was also used to calculate three additional variables: average number of eggs per female, average number of juveniles per female, and average number of offspring per female ((# eggs + # juveniles)/# females). These three ratios were then used to approximate fertilization and maturity rates between Weeks 3 and 4. Juvenile sizes were also obtained by using the freehand trace tool and measuring the number of pixels encapsulated. Sizes were then converted to |m^2^ using a conversion factor of 71330 pixels per 62,500|m^2^, which was calculated by measuring the area of a photo of a 0.0625mm^2^ hemocytometer cell at 40x magnification.

### Statistical Analysis

All count outcome variables were analyzed using linear mixed models with temperature, pH, and their interaction as fixed effects and Dish ID as a random effect. In order to meet the parametric assumptions of normality of residuals and homogeneity of variances, all count data was subjected to a square-root transformation as needed before being analyzed. We tested the significance of our fixed effects by conducting log-likelihood tests via model comparison using the anova function, where one model included the effect of interest while the other model excluded it.

We analyzed the proportion of productive females using a generalized linear mixed model with a beta distribution. Size data was also analyzed with a generalized linear mixed model using a gamma distribution. GLMMs included temperature, pH, and their interactions as fixed effects, and petri dish ID as a random effect. Average number of gametophytes per photo in a given dish was also calculated and included in the size model as a covariate to account for possible density dependence. We also separately analyzed the relationship between average size of juveniles per photo and the covariate (average number of gametophytes per photo) using a linear regression model that included only the covariate as a fixed effect.

All count and size data were only analyzed for Week 4 of our experiment, but calculated ratios of eggs per female (eggs/fem), juveniles per female (juvs/fem), and offspring per female (offspring/fem) were analyzed for both Weeks 3 and 4 in order to draw conclusions about differences in rates of fertilization or maturation. Specifically, we used the ratio of offspring/fem to ask whether females, regardless of treatment, were making the same effort to reproduce, and the ratios of eggs/fem and juvs/fem were calculated to inform us about which stage reproduction was within each treatment. We tested the significance of our fixed effects via model comparison using the anova function, where one model included the effect of interest while the other model excluded it. Hypothesis testing was conducted via log-likelihood tests for count and offspring ratio LMMs and chi-squared tests for size GLMMs. All analyses were performed using R version 4.1.2 (R Core Team 2021) and the packages *nlme* (Pinheiro et al. 2022), *lme4* (Bates et al. 2015), and *glmmTMB* (Brooks et al. 2017).

## Results

### Female and Male Gametophyte Development

High temperatures resulted in a significant decline in the number of females present in Week 4 (Log-Likelihood=51.1283, DF=36, p<0.0001) (Figure 2, Table 1) . Neither pH nor the interaction between pH and temperature had a significant effect on female gametophyte numbers. The random effect of Dish had a significant effect on female gametophyte numbers in Week 4 (Log-Likelihood=7.8795, p=0.005), indicating that there was some variation between petri dishes for this test.

**Figure 2:**
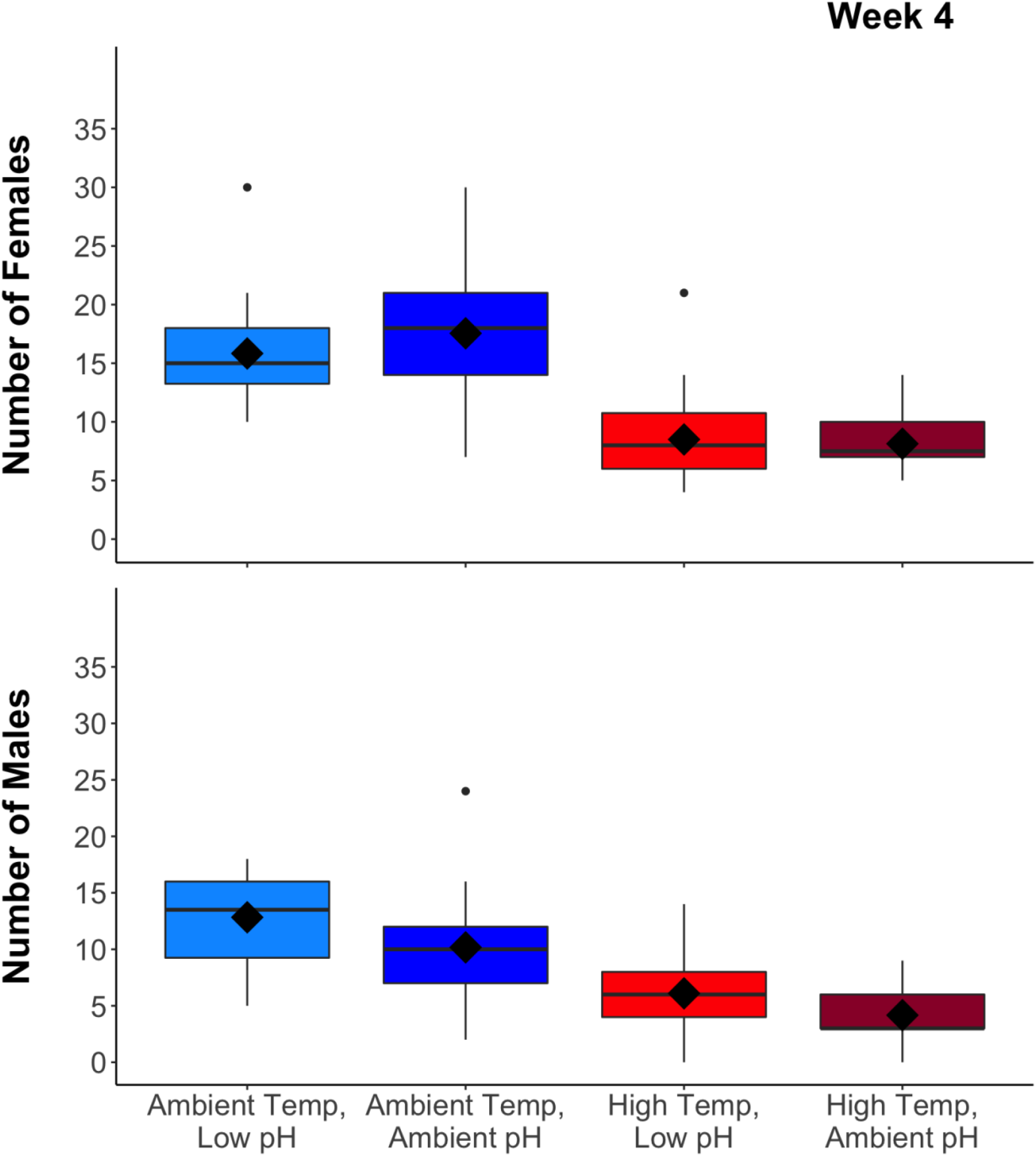
Female and male gametophytes present in each photo after 4 weeks of growth. The box plots summarize the mean (diamond) and median (box midline) for each treatment, the first and third quartiles (upper and lower box limits), outliers within 1.5 times the inter-quartile range (vertical lines), and outliers beyond that range (dots).

**Table 1:** Means and SDs of Count data. Counts represent the mean value of three photos per 10 dishes in each treatment.

| **Treatment** | **Eggs** | **Females** | **Juveniles** | **Males** |
| --- | --- | --- | --- | --- |
| Ambient Temp, Low pH | 15.6 ± 10.48 | 15.83 ± 4.05 | 20.03 ± 6.99 | 12.83 ± 3.79 |
| Ambient Temp, Ambient pH | 12.24 ± 6.65 | 17.55 ± 5.13 | 21.72 ± 6.14 | 10.17 ± 4.26 |
| High Temp, Low pH | 6.03 ± 5.17 | 8.5 ± 3.88 | 12 ± 5.26 | 6.1 ± 3.17 |
| High Temp, Ambient pH | 3.17 ± 2.74 | 8.13 ± 2.21 | 12.47 ± 4.29 | 4.17 ± 2.29 |

Males were similarly negatively affected by high temperatures in both Week 3 (Log-Likelihood=18.9288, DF=36, p<0.0001) and Week 4 (Log-Likelihood=45.393, DF=36, p<0.0001). However, they varied from females in that their numbers were significantly higher under lower pH conditions in Week 4 (Log-Likelihood=8.6378, DF=36, p=0.0033). Neither the pH:temperature interaction term nor the random effect Dish had any significant effect on male gametophyte numbers. In summary, these results indicate that temperature caused a significant decrease in female and male gametophyte numbers, whereas low pH only caused a significant increase in male gametophyte numbers.

### Egg and Sporeling Counts

After 4 weeks, high temperatures were correlated with significantly lower numbers of both eggs (Log-Likelihood=33.73, DF=36, p<0.0001) and juveniles (Log-Likelihood=36.6391, DF=36, p<0.0001) (Figure 3). Low pH was associated with significantly larger numbers of eggs (Log-Likelihood=4.3958, DF=36, p=0.036) whereas there were no significant effects on juvenile counts (Log-Likelihood=1.0702, DF=36, p=0.3009). The interaction term for pH:temperature was insignificant for counts of both eggs and juveniles. The random effect of Dish was insignificant for juveniles in Week 4 (Log-Likelihood=3.0544, p=0.0805), but was significant for eggs in Week 4 (Log-Likelihood=5.2080, p=0.0225). Overall, temperature caused the greatest decreases in both egg and juvenile numbers, whereas low pH caused a significant increase in eggs only.

**Figure 3:**
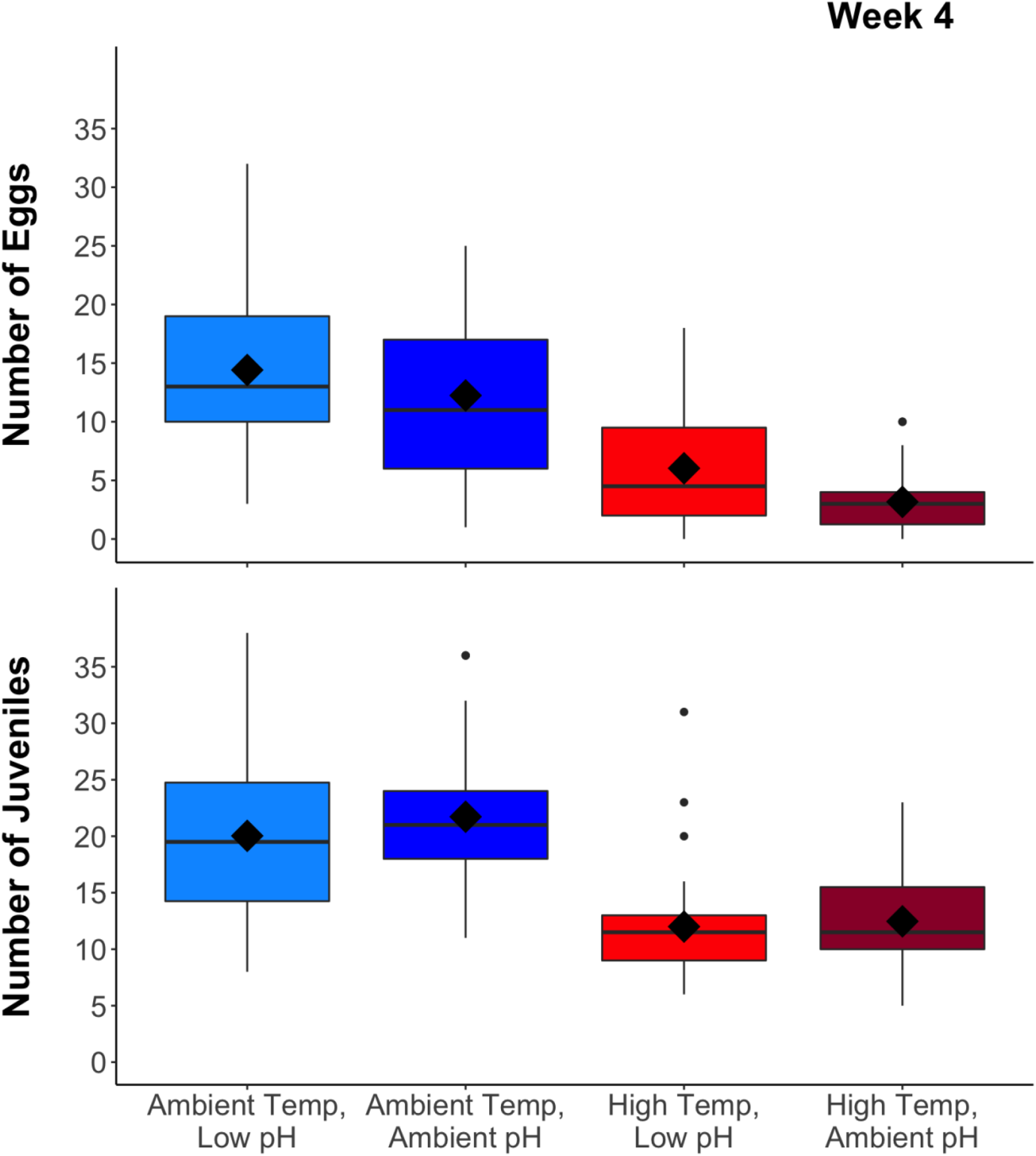
Eggs and juveniles present in each photo after 4 weeks of growth. The box plots summarize the mean (diamond) and median (box midline) for each treatment, the first and third quartiles (upper and lower box limits), outliers within 1.5 times the inter-quartile range (vertical lines), and outliers beyond that range (dots).

### Proportion of Productive Females

The proportion of productive females was uniformly high across all treatments, but the high temperature treatments consistently resulted in nearly 100% of females reaching productivity by Week 4 (Figure 4). We found that the proportion of productive females was not significantly affected by the interaction between temperature and pH (Chi-Sq=0.5117, p=0.4744) nor the individual effect of pH (Chi-Sq=1.1619, p=0.2811). High temperature was the only variable to result in a significant increase in the proportion of productive females (Chi-Sq=28.187, p<0.0001).

**Figure 4:**
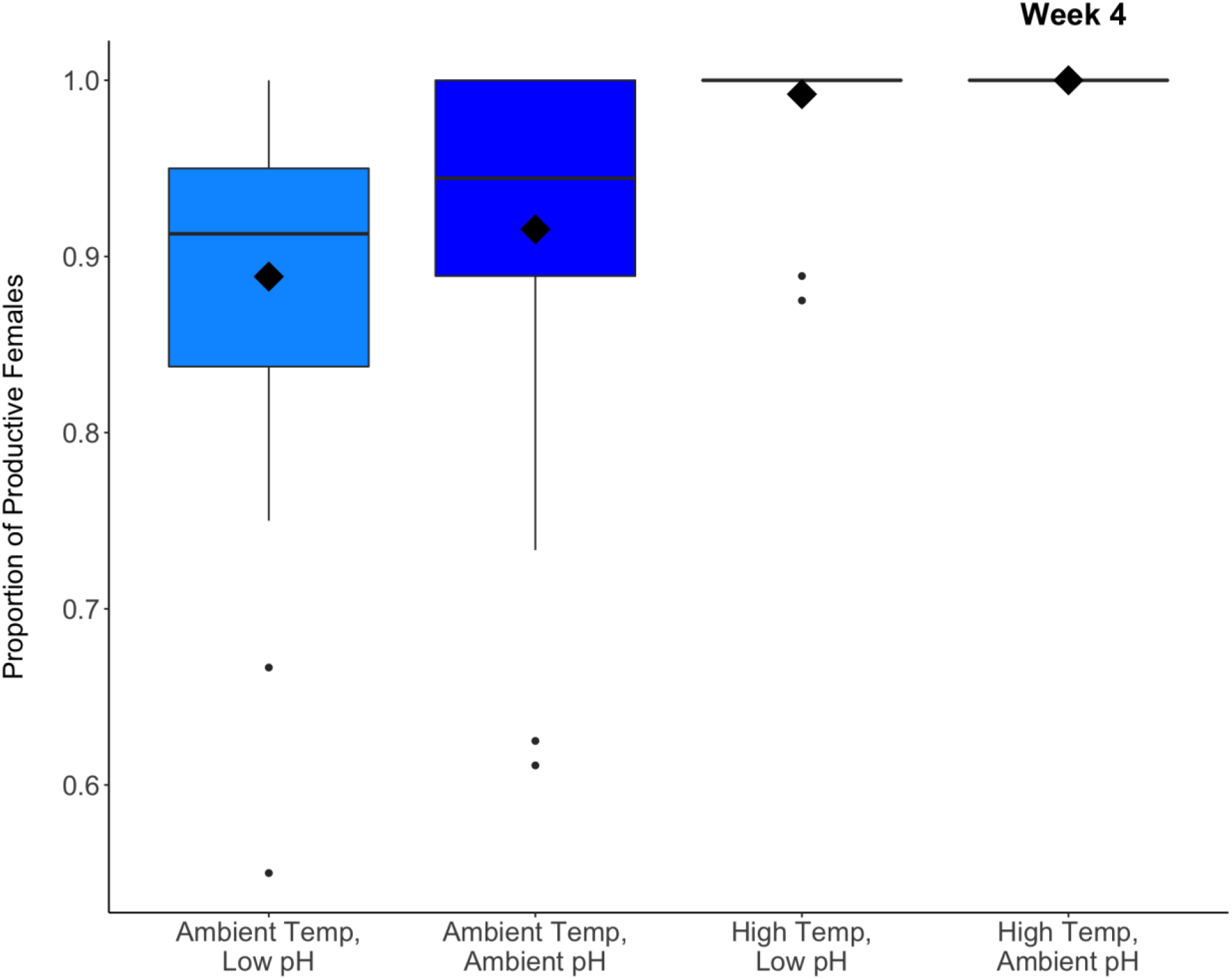
Proportion of productive female gametophytes after 4 weeks of growth. The box plots summarize the mean (diamond) and median (box midline) for each treatment, the first and third quartiles (upper and lower box limits), outliers within 1.5 times the inter-quartile range (vertical lines), and outliers beyond that range (dots).

### Ratios of Offspring per Female

For mean number of eggs per female (egg/fem), we found a marginally significant effect of the interaction between temperature and pH in Week 3 (Log-Likelihood=3.7737, DF=36, p=0.0521) but not Week 4 (Log-Likelihood=0.0976, DF=36, p=0.7547) (Table 2, Figure 5). Investigating temperature and pH individually in Week 3, we found that low pH (Log-Likelihood=3.7345, DF=36, p=0.0533) was marginally significantly associated with a decreased egg/fem ratio under ambient temperature treatments, but an increased egg/fem ratio under high temperature treatments. Temperature was found to be insignificant in Week 3 (Log-Likelihood=0.1406, DF=36, p=0.7077). In Week 4, low pH was found to be significantly associated with a higher egg/fem (Log-Likelihood=9.3663, DF=36, p=0.0022) whereas low temperature resulted in lower egg/fem (Log-Likelihood=13.114, DF=36, p=0.0003). The effect of Dish was insignificant in both Week 3 (Log-Likelihood=0.2318, p=0.6302) and Week 4 (Log-Likelihood=0.5643, p=0.4525).

**Figure 5:**
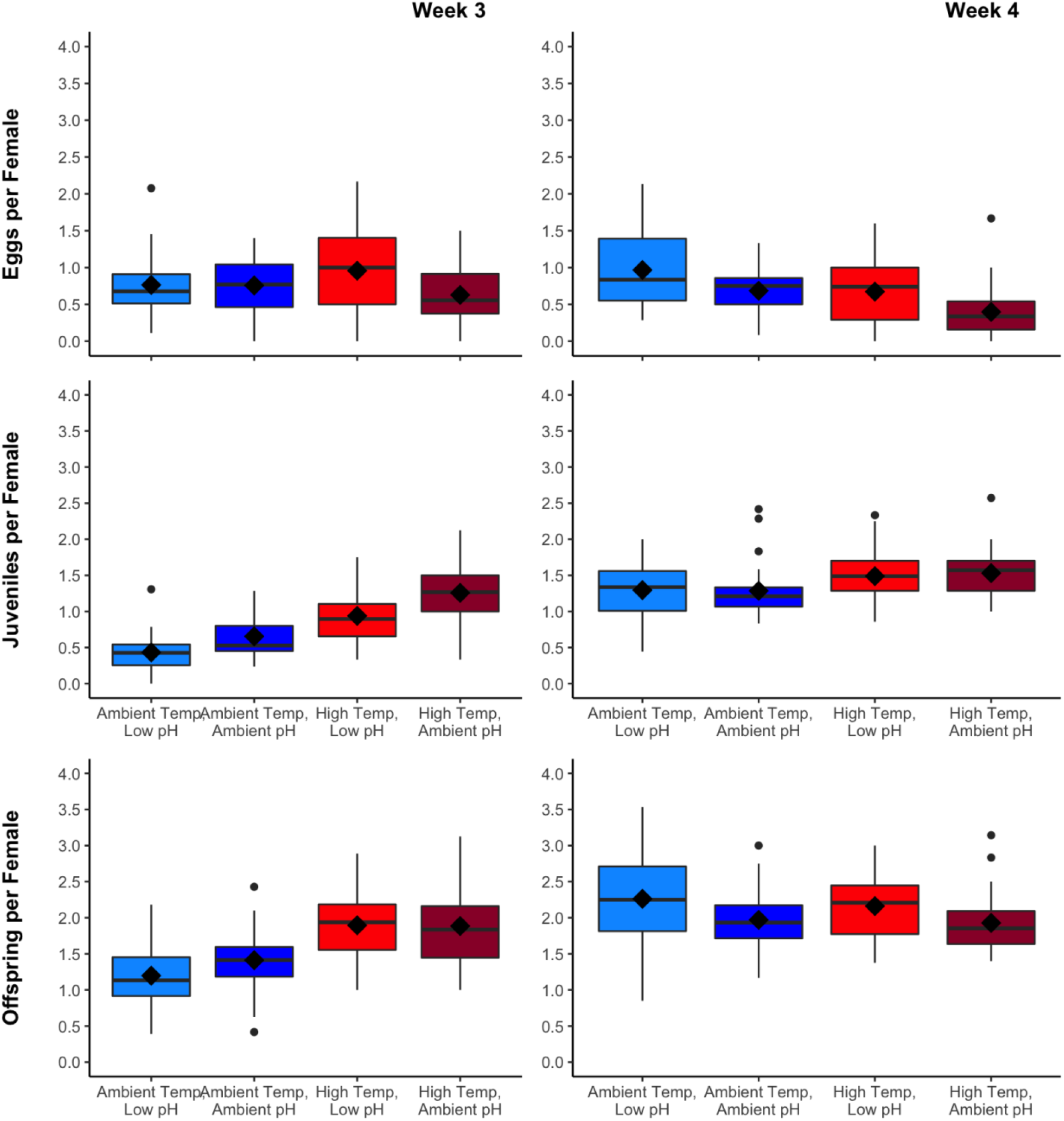
Eggs, juveniles, and total offspring (eggs + juveniles) per female after 3 and 4 weeks of growth. The box plots summarize the mean (diamond) and median (box midline) for each treatment, the first and third quartiles (upper and lower box limits), outliers within 1.5 times the inter-quartile range (vertical lines), and outliers beyond that range (dots).

**Table 2:** Means and SDs of Productivity Data. The mean productivity is the number of offspring per female per three photos per 10 dishes in each treatment.

**Week 3**
| **Treatment** | **Proportion Productive Females** | **Eggs per Female** | **Juveniles per Female** | **Total Offspring per Female** |
| --- | --- | --- | --- | --- |
| Ambient Temp, Low pH | 0.68 ± 0.2 | 0.76 ± 0.4 | 0.43 ± 0.27 | 1.2 ± 0.48 |
| Ambient Temp, Ambient pH | 0.8 ± 0.15 | 0.76 ± 0.4 | 0.66 ± 0.29 | 1.41 ± 0.41 |
| High Temp, Low pH | 0.89 ± 0.13 | 0.96 ± 0.55 | 0.94 ± 0.35 | 1.9 ± 0.49 |
| High Temp, Ambient pH | 0.95 ± 0.08 | 0.63 ± 0.42 | 1.26 ± 0.42 | 1.89 ± 0.53 |

| **Treatment** | **Proportion Productive Females** | **Eggs per Female** | **Juveniles per Female** | **Total Offspring per Female** |
| --- | --- | --- | --- | --- |
| Ambient Temp, Low pH | 0.89 ± 0.11 | 0.97 ± 0.53 | 1.3 ± 0.4 | 2.26 ± 0.65 |
| Ambient Temp, Ambient pH | 0.92 ± 0.1 | 0.68 ± 0.3 | 1.29 ± 0.37 | 1.97 ± 0.42 |
| High Temp, Low pH | 0.99 ± 0.03 | 0.67 ± 0.48 | 1.49 ± 0.38 | 2.16 ± 0.49 |
| High Temp, Ambient pH | 1 ± 0 | 0.4 ± 0.36 | 1.53 ± 0.33 | 1.93 ± 0.41 |

**Table 3:**
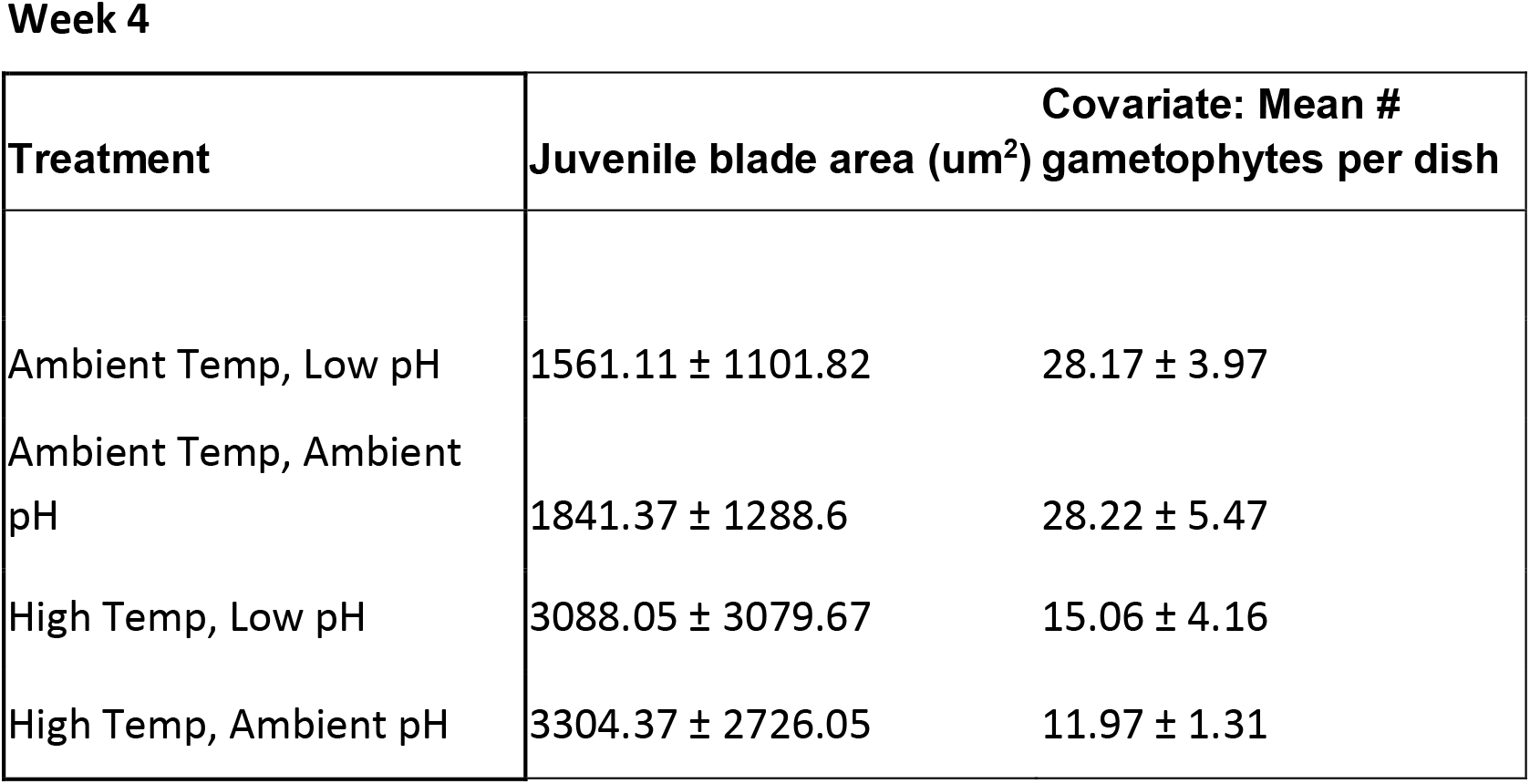
Means and SDs of sizes and covariates. Size represents the area of juvenile blades in um2, where as the covariate refers to the mean number of gametophytes per dish.

For mean number of juveniles per female (juv/fem), we found that high temperatures resulted in larger juv/fem in both Week 3 (Log-Likelihood=45.2639, DF=36, p<0.0001) and Week 4 (Log-Likelihood=9.1867, DF=36, p=0.0024). Low pH treatments had significantly lower juv/fem ratio than ambient pH treatments in Week 3 (Log-Likelihood=16.7485, DF=36, p<0.0001) but not in Week 4 (Log-Likelihood=0.1262, DF=36, p=0.7225). Neither the interaction term pH:temperature nor the random effect Dish were significant in either week. Increased ratios of total offspring per female (offspring/fem) were significantly correlated with high temperatures in Week 3 (Log-Likelihood=30.4186, DF=36, p<0.0001) but not Week 4 (Log-Likelihood=0.2279, DF=36, p=0.6331), whereas more offspring/fem were significantly correlated with low pH in Week 4 (Log-Likelihood=5.2345, DF=36, p=0.0221) but not Week 3 (Log-Likelihood=1.3622, DF=36, p=0.2432). Neither the interaction between temperature and pH nor the random effect Dish were significant in either week.

Across all ratios, high temperature had the greatest impacts in Week 3, resulting in lower ratios of juvs/fem and offspring/fem, whereas low pH was most significant in Week 4, resulting in high eggs/fem and offspring/fem.

### Growth of Juvenile Bull Kelp

When analyzing the global trend across all treatments, we found that juvenile size was significantly influenced by the covariate (average number of gametophytes within each dish) in Week 4 (R^2^ =0.639, p<0.0001), indicating possible density dependence where increased number of gametophytes resulted in significantly smaller juveniles (Figure 6). When included in the GLMM, the 3-way interaction between pH, temperature, and the covariate was significant (Chi-Sq=6.3387, p=0.0118), but all 2-way interactions were insignificant.

**Figure 6:**
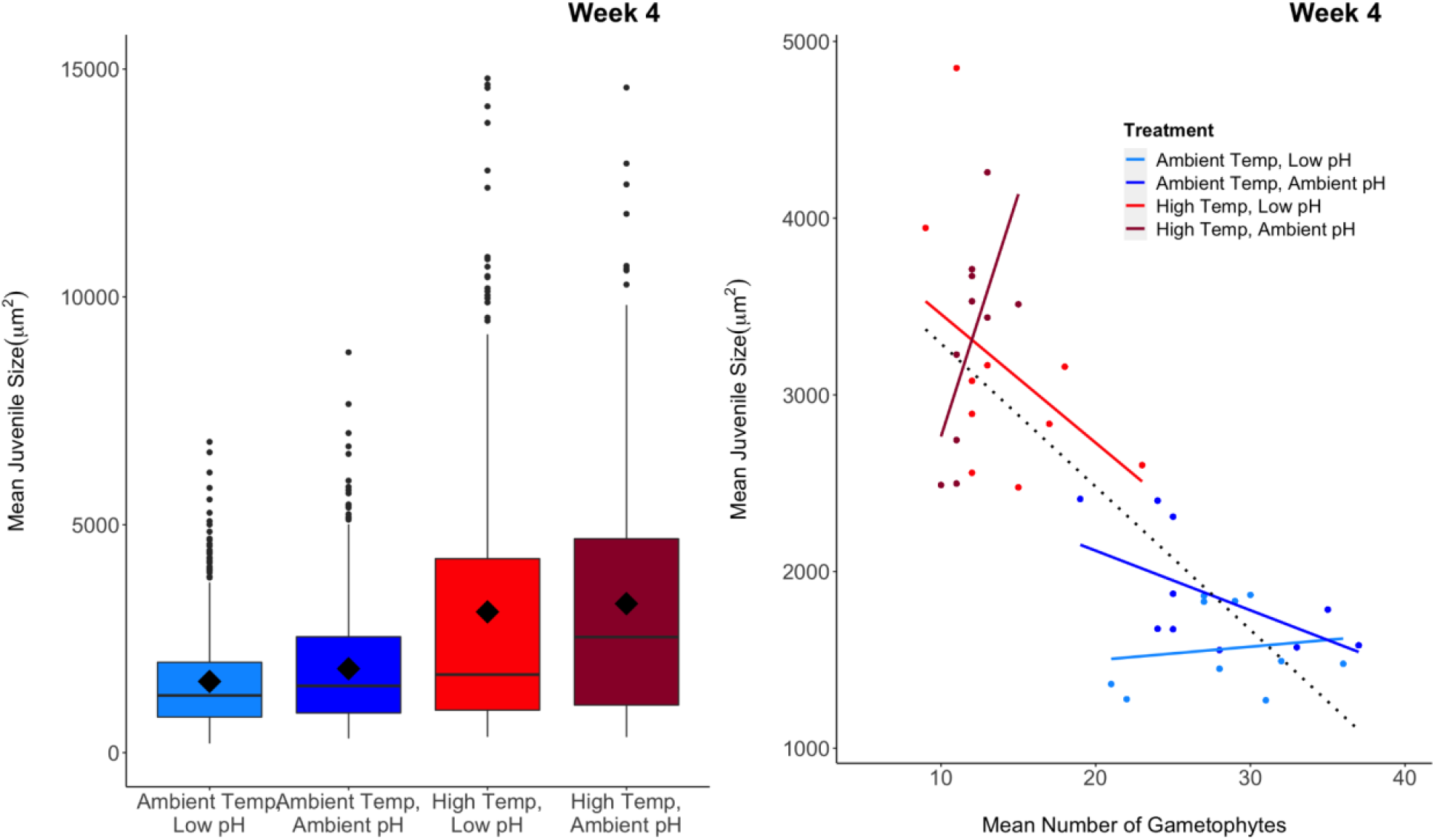
Left panel shows the average size of juveniles after 4 weeks of growth. The left panel shows box plots that summarize the mean (diamond) and median (box midline) for each treatment, the first and third quartiles (upper and lower box limits), outliers within 1.5 times the inter-quartile range (vertical lines), and outliers beyond that range (dots). The right panel shows the relationship of the covariate (mean number of gametophytes) to the response variable (mean juvenile size). Trends within groups are represented by colored lines, whereas the trend across all groups is represented by the dotted black line. Heterogeneous slopes and different ranges of values for each treatment indicate that the different treatments are confounded with differences in the covariate.

In order to examine the role of each fixed factor (pH, temperature, and the covariate) in the 3-way interaction, we subset our model within the two pH and two temperature levels to elucidate the significance of the covariate and the other non-subset factor. When low pH and ambient pH treatments were separately analyzed, the covariate was never independently significant, but high temperatures resulted in a significant increase in size by itself under low pH conditions (Chi-Sq=10.051, p=0.0015) and the temperature by covariate interaction was significant under ambient pH conditions (Chi-Sq=5.3001, p=0.02132). When the two temperature treatments were analyzed separately, the covariate was again never significant by itself, but pH resulted in a significant decrease in juvenile sizes under ambient temperature conditions (Chi-Sq=3.955, p=0.04673), and the pH by covariate interaction had a significant effect on size under high temperature conditions (Chi-Sq=6.0391, p=0.01399) (Figure 6). In summary, juvenile sizes were most significantly impacted by high temperatures, but pH and the number of gametophytes present greatly influenced this effect.

## Discussion

Our results demonstrated that both temperature and pH do significantly impact bull kelp reproduction and development, but the effects were more variable and nuanced than we predicted. Our most consistent finding was that high temperatures resulted in significant reduction in the number of gametophytes that survived and/or settled and the number of eggs and juveniles produced. The number of male gametophytes, female gametophytes, eggs, and juveniles were always lower in high temperature treatments than ambient temperature treatments, regardless of pH. These results align with previous findings that bull kelp exposed to increased temperature conditions had reduced spore germination rates and reduced gametophyte growth (Lind and Konar 2017; Muth et al 2019, Schiltroth 2021). Increased temperatures also result in decreased gametophyte survival and sporophyte recruitment in numerous other kelp species including giant kelp (*Macrocystis pyrifera*) (Hollarsmith et al. 2020), stalked kelp (*Pterygophora californica*) (Howard 2014), spiny kelp (*Ecklonia radiat*a) (Alsuwaiyan 2021), sugar kelp (*Saccharina latissima*), and dragon kelp (*Eualaria fistulosa*) (Lind and Konar 2017).

Low pH had a non-significant and lower effect size on the number of female gametophytes or juveniles than temperature in our study, but there was a significant increase in male gametophytes and eggs. Other studies have found varying impacts of pH on kelp gametophyte growth and survival. Several studies have found overall positive effects of low pH on *M. pyrifera* gametophyte growth, survival, and size (Roleda et al. 2012, Leal et al. 2017), where as other studies found that elevated *p*CO2 had little effect on rates of growth and photosynthesis (Fernández et al. 2015) or reproduction (Hollarsmith et al 2020). The variation in kelp organismal responses across studies, species, and location (Roleda and Hurd 2012; Hollarsmith et al 2020) indicates that there is much more to be understood about the impacts of ocean acidification on kelp reproduction.

Our results also support a hypothesis that high temperatures may result in bull kelp gametophytes potentially allocating more effort into reproduction. While the overall number of gametophytes and juveniles declined under high temperature conditions, the female gametophytes that survived were more productive on average, and produced more juveniles earlier than female gametophytes under ambient temperature conditions. This result suggests that female gametophytes possessed similar capacities for offspring production regardless of treatment, but that females become reproductive sooner under high temperature conditions. This idea is also supported by the separate ratios for eggs per female and juveniles per female in each week.

We also saw that our results align with this hypothesis via slower reproduction and development under ambient temperature conditions. In Week 3, high temperature treatments had higher ratios of both eggs/fem and juvs/fem than ambient temperature treatments. By Week 4, however, the eggs/fem ratio in ambient temperature treatments exceeded that of high temperature treatments, and the juvs/fem ratio was similar regardless of temperature treatment. The later increases in egg/fem and juvs/fem ratios in ambient temperatures seem to indicate that female gametophytes have equal reproductive capacity under both our temperature treatments, but females growing under high temperature treatments were progressing through reproduction earlier. These results are consistent with those of Howard (2014), who found that ambient temperatures lead to slower microstage reproduction of bull kelp as well as giant kelp (*Macrocystis pyrifera*) and stalked kelp (*Pterygophora californica*)..

In contrast, low pH conditions seemed to impact reproductive efforts differently based on temperature conditions. Under ambient temperatures, low pH appeared to slow reproduction by producing the lowest egg/fem, juv/fem, and offspring/fem ratios of all treatments in Week 3, but then increased enough to match the ratios of other treatments in Week 4 (Figure 5). Under high temperature conditions on the other hand, low pH treatments produced the highest egg/fem and offspring/fem ratios and second highest juv/fem ratios of all treatments in Week 3, with only nominal increases in these ratios in Week 4. Additionally, the lowest proportion of productive females was observed in low pH treatments under ambient temperatures (Figure 4), and these females seemed to produce more offspring per female, later in the experiment. The greatest increase in productivity between Week 3 and Week 4 under any pH treatments can be found in the large increase in egg/fem ratios under ambient temperature conditions. This late increase in production of eggs may potentially signal that a delay in reproduction occurs under low pH and ambient temperature conditions.

Recent advances in kelp reproduction studies have given needed attention to delayed development of microscopic stages and the resulting “bank of microscopic forms” (Hoffman and Santelices 1991, Carney and Edwards 2006, Schoenrock et al. 2021), but less focus has been placed on the factors that may accelerate microscopic kelp development. In terrestrial plants, increased temperatures have been found to result in an acceleration of pollen tube growth and stigma and ovule development, which correspond to an overall reduction of the length of time females are receptive to pollination (Hedhly et al. 2009). Reviews of other marine organisms, specifically benthic invertebrates, have shown that increased sea surface temperatures may increase the rate and timing of development and spawning (Przeslawski et al. 2008). In order to better understand the ability of populations to recover from extreme climate disturbance events, more research is needed to better understand the effect of climate stressors on survival, time to development, and propagule production.

For all variables other than egg/fem in Week 3, we found no significant temperature and pH interactions on bull kelp microscopic stages, which contrasts with studies of giant kelp, that did find interactive effects of pH or *p*CO2 and temperature on spore, gametophyte, and sporeling production (Gaitan-Espitia et al. 2014, Shukla and Edwards 2017, Hollarsmith et al. 2020). Our results interestingly reflect natural seasonal fluctuations in northern California’s coastal waters, where conditions of low pH and high temperature rarely overlap. Oceanographic conditions in California’s coastal waters are typically determined by three seasons: upwelling season (April to June), relaxation season (July to October), and storm season (October to March) (García-Reyes and Largier 2012). Upwelling season is characterized by the upwelling of cold, dense, nutrient rich water from the deep ocean to the coastline, causing coastal waters to be colder and more acidic. During non-upwelling season, on the other hand, the intensity of upwelling processes decreases, and coastal waters become relatively warmer, more basic, and exhibit less primary productivity and chlorophyll-a content due to shortened day lengths and decreased nutrient concentrations (García-Reyes and Largier 2012). The bull kelp lifecycle has a relatively consistent, yet still flexible, phenology. The majority of visible juveniles appear in the spring (upwelling season) and most adults become reproductive by the end of July (relaxation season), but these two events of visible recruitment and spore release have been observed to occur in all seasons, albeit at much lower rates (Maxell and Miller 1996, Dobkowski et al. 2018).

As a result, the need for gametophytes to endure high temperature or low pH conditions is dependent on the life cycle of the parent sporophyte. Gametophytes and sporelings that develop in the spring will likely be most exposed to low temperatures and low pH as a result of upwelling, but the vast majority of gametophytes and sporelings that develop in the fall will be exposed to non-upwelling season late summer conditions. As such, it is conceivable that high temperatures in September and October (relaxation season) would affect the first month of juvenile development, whereas upwelling season would likely be more important for visible juveniles than the microscopic ones.

However, our results potentially contrast with those of Dobkowski et al. 2018 in that we found that low pH (most often seen in the Spring upwelling season) resulted in slower reproduction and growth whereas high temperature (most often seen in late summer and fall) accelerated it. In their study, Dobkowski et al. (2018) witnessed the quickest recruitment of visible juveniles (indicating faster microscopic development times) in the spring (upwelling season), and slowest recruitment (implying slower microscopic development times) in the late summer and fall (relaxation season. A potential explanation for the different observed reproductive rates is that Dobkowski et al. conducted their experiments in the field, where they were exposed to a full of array of abiotic conditions, whereas our experiments were conducted in a laboratory setting where only temperature and pH were manipulated, and all other variables were held constant, including nutrients. Previous studies have shown that delayed development of microscopic kelp stages is often closely tied to insufficient nutrient and light regimes (Carney and Edwards 2010), both of which are present between September to March due to dampened upwelling conditions and reduced daylength. As such, the slow development over winter in natural populations suggests that nutrients from upwelling and daylength could be more important than temperature and pH fluctuations in promoting the development of microscopic kelp stages.

Our results suggest that there may be some density dependence effects on juvenile growth at these microscopic stages. The difference in juvenile sizes between treatments was most significantly correlated with temperature, but also showed at least a marginally significant correlation with the number of gametophytes present in both weeks (Figure 5). However, due to the fact that high temperatures consistently resulted in significant decreases in gametophyte numbers, the relationships of both temperature and number of gametophytes to gametophyte size are confounded, and direct causation cannot be determined. As a result, more research is needed to see whether these increased sizes were really a result of high temperatures or whether they were a result of lowered density of individuals.

In natural populations, there are numerous density-dependent effects that impact kelp reproduction and recruitment. At initial spore settlement, high densities of gametophytes are needed for fertilization between male and female gametophytes to occur, so Allee effects may occur if spores settle at a density of less than 1 spore/mm^2^ (Reed 1990). The direction of density-dependence then reverses somewhere between the gametophyte stage and the point where a juvenile becomes easily detectable to the naked eye, and numerous kelps, including bull kelp, exhibit subsequent increases in mortality until they reach the adult life stage (Schiel and Foster 2006, Reed et al. 1991, Dobkowski et al. 2018). Our petri dishes, each containing thousands of spores and gametophytes, contained a bottom surface area of approximately 157.8cm^2^, whereas an adult bull kelp holdfast can grow up to an estimated 40cm in diameter and 1256cm^2^ surface area (Abbot and Hollenberg 1976). As a result, less than one juvenile sporophyte in each petri dish would be expected to survive to adulthood and reproductive maturity in natural environments.

The results of this research indicate that climate change will significantly affect bull kelp reproduction via increased temperatures, and, to a lesser extent, ocean acidification. Increasing frequency and intensity of extreme temperature events such as marine heatwaves will likely lead to a massive decrease in the survival of gametophytes and decreased, but accelerated, production of juvenile sporophytes. Lowered pH, mimicking ocean acidification, resulted in an increase in numbers of male gametophytes and juvenile sporelings, as well as a slower reproduction rate.

The ability of bull kelp to recover from extreme climate events depends on the ability of all lifestages to withstand abiotic stress. Currently, managers and scientists are focused on understanding the natural processes that guide kelp recovery and whether those may require some degree of management intervention. In order to better understand the ability of species to recover from climate change, more research is needed to understand the ability of intermediate stages between propagules and adults to withstand abiotic stress. In order to intervene successfully through restoration, an understanding of potential bottlenecks and challenges present at each life stage is necessary. This study lends a little insight into the otherwise largely unknown world of microscopic life stages of kelp

## Acknowledgments

We would like to thank Rob Coyan for assistance collecting sporophylls, and Carol Vines for help with microscopy methods. A. Blandino of the UC Davis Statistical Consulting Group and K. Laskowksi provided valuable statistical advice for this manuscript. Laboratory experiments were conducted at Bodega Marine Lab (BML), and were aided greatly by the assistance of many BML staff members.

## SUPPLEMENTARY TABLES

**Table 1:**
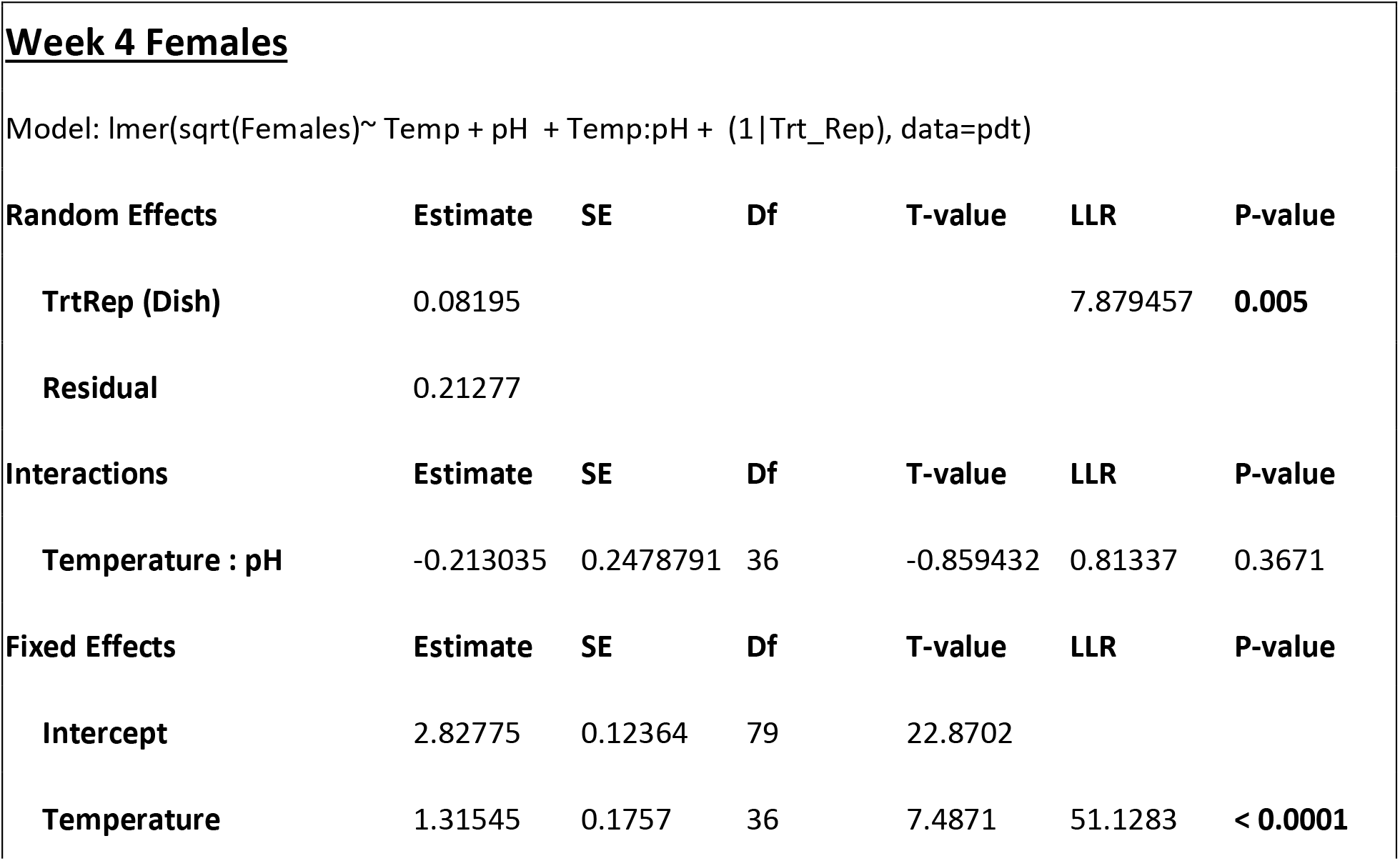
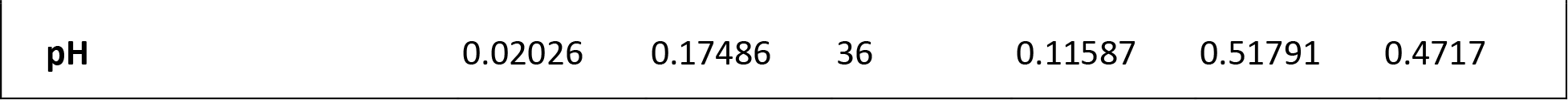
Statistical outcomes for models of female count data.

| **<u>Week 4 Females</u>** |  |  |  |  |  |  |
| --- | --- | --- | --- | --- | --- | --- |
| Model: lmer(sqrt(Females)~ Temp + pH + Temp:pH + (1\|Trt_Rep), data=pdt) |  |  |  |  |  |  |
| **Random Effects** | **Estimate** | **SE** | **Df** | **T-value** | **LLR** | **P-value** |
| **TrtRep (Dish)** | 0.08195 |  |  |  | 7.879457 | **0.005** |
| **Residual** | 0.21277 |  |  |  |  |  |
| **Interactions** | **Estimate** | **SE** | **Df** | **T-value** | **LLR** | **P-value** |
| **Temperature : pH** | -0.213035 | 0.2478791 | 36 | -0.859432 | 0.81337 | 0.3671 |
| **Fixed Effects** | **Estimate** | **SE** | **Df** | **T-value** | **LLR** | **P-value** |
| **Intercept** | 2.82775 | 0.12364 | 79 | 22.8702 |  |  |
| **Temperature** | 1.31545 | 0.1757 | 36 | 7.4871 | 51.1283 | **< 0.0001** |
| **pH** | 0.02026 | 0.17486 | 36 | 0.11587 | 0.51791 | 0.4717 |

**Table 2:**
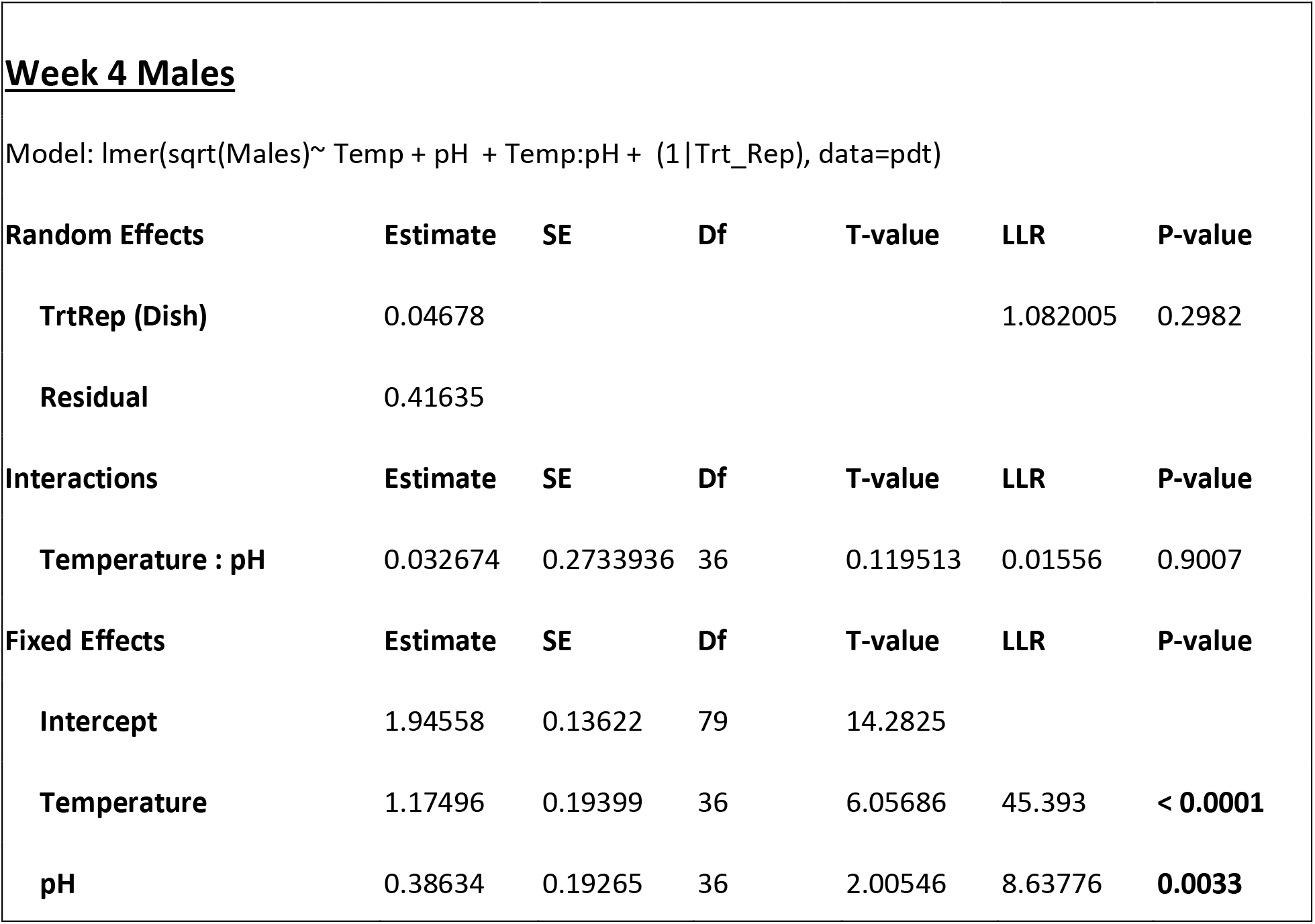
Statistical outcomes for models of male count data.

|  |  |  |  |  |  |  |
| --- | --- | --- | --- | --- | --- | --- |
| **<u>Week 4 Males</u>** |  |  |  |  |  |  |
| Model: lmer(sqrt(Males)~ Temp + pH + Temp:pH + (1\|Trt_Rep), data=pdt) |  |  |  |  |  |  |
| **Random Effects** | **Estimate** | **SE** | **Df** | **T-value** | **LLR** | **P-value** |
| TrtRep (Dish) | 0.04678 |  |  |  | 1.082005 | 0.2982 |
| Residual | 0.41635 |  |  |  |  |  |
| **Interactions** | **Estimate** | **SE** | **Df** | **T-value** | **LLR** | **P-value** |
| Temperature : pH | 0.032674 | 0.2733936 | 36 | 0.119513 | 0.01556 | 0.9007 |
| **Fixed Effects** | **Estimate** | **SE** | **Df** | **T-value** | **LLR** | **P-value** |
| Intercept | 1.94558 | 0.13622 | 79 | 14.2825 |  |  |
| Temperature | 1.17496 | 0.19399 | 36 | 6.05686 | 45.393 | < 0.0001 |
| pH | 0.38634 | 0.19265 | 36 | 2.00546 | 8.63776 | 0.0033 |

**Table 3:**
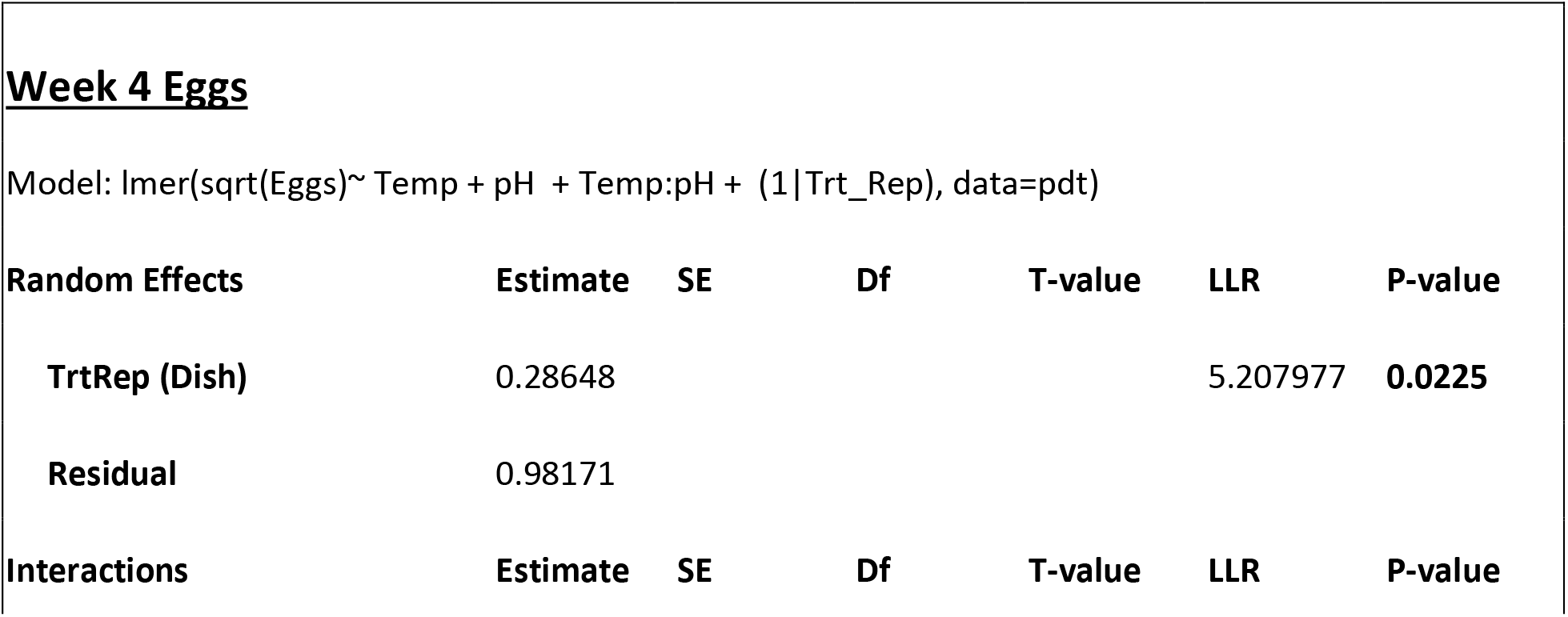
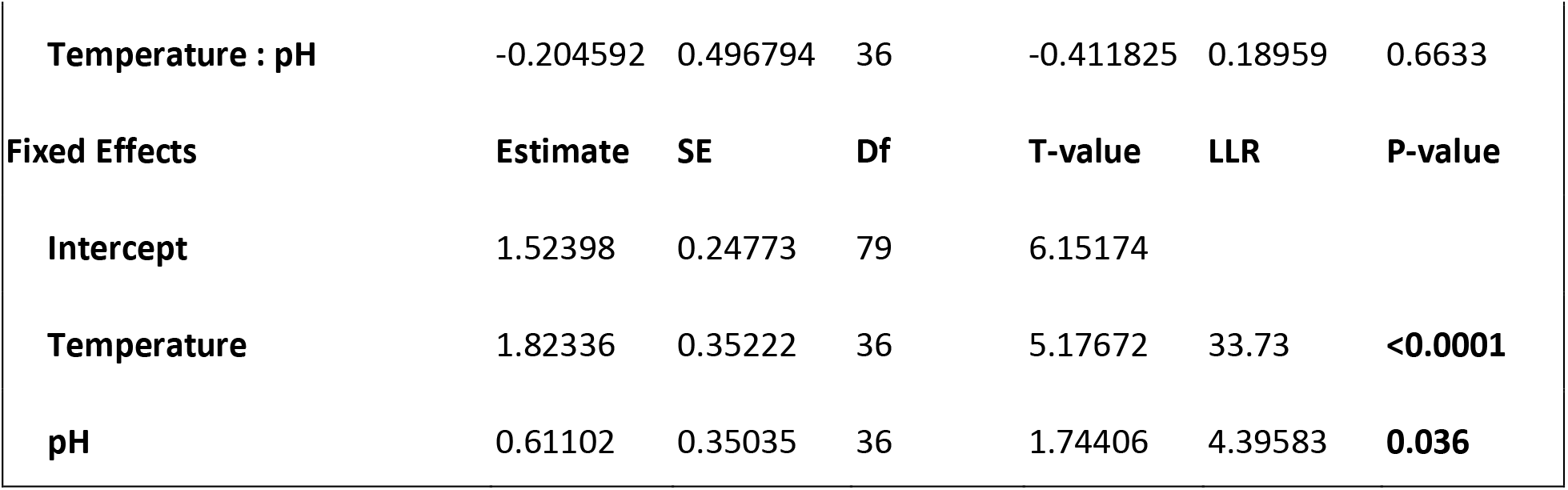
Statistical outcomes for models of egg count data.

|  |  |  |  |  |  |  |
| --- | --- | --- | --- | --- | --- | --- |
| **<u>Week 4 Eggs</u>** |  |  |  |  |  |  |
| Model: lmer(sqrt(Eggs)~ Temp + pH + Temp:pH + (1\|Trt_Rep), data=pdt) |  |  |  |  |  |  |
| **Random Effects** | **Estimate** | **SE** | **Df** | **T-value** | **LLR** | **P-value** |
| TrtRep (Dish) | 0.28648 |  |  |  | 5.207977 | 0.0225 |
| Residual | 0.98171 |  |  |  |  |  |
| **Interactions** | **Estimate** | **SE** | **Df** | **T-value** | **LLR** | **P-value** |
| **Temperature : pH** | -0.204592 | 0.496794 | 36 | -0.411825 | 0.18959 | 0.6633 |
| **Fixed Effects** | **Estimate** | **SE** | **Df** | **T-value** | **LLR** | **P-value** |
| **Intercept** | 1.52398 | 0.24773 | 79 | 6.15174 |  |  |
| **Temperature** | 1.82336 | 0.35222 | 36 | 5.17672 | 33.73 | **<0.0001** |
| **pH** | 0.61102 | 0.35035 | 36 | 1.74406 | 4.39583 | **0.036** |

**Table 4:** Statistical outcomes for models of juvenile count data.

|  |  |  |  |  |  |  |
| --- | --- | --- | --- | --- | --- | --- |
| **<u>Week 4 Juveniles</u>** |  |  |  |  |  |  |
| Model: lmer(sqrt(Juveniles)~ Temp + pH + Temp:pH + (1\|Trt_Rep), data=pdt) |  |  |  |  |  |  |
| **Random Effects** | **Estimate** | **SE** | **Df** | **T-value** | **LLR** | **P-value** |
| **TrtRep (Dish)** | 0.08057 |  |  |  | 3.054386 | 0.0805 |
| **Residual** | 0.38809 |  |  |  |  |  |
| **Interactions** | **Estimate** | **SE** | **Df** | **T-value** | **LLR** | **P-value** |
| **Temperature : pH** | -0.122581 | 0.290655 | 36 | -0.42174 | 0.19862 | 0.6558 |
| **Fixed Effects** | **Estimate** | **SE** | **Df** | **T-value** | **LLR** | **P-value** |
| **Intercept** | 3.48191 | 0.14489 | 79 | 24.0311 |  |  |
| **Temperature** | 1.13264 | 0.20614 | 36 | 5.49452 | 36.6391 | **<0.0001** |
| **pH** | -0.0829 | 0.20491 | 36 | -0.4047 | 1.07023 | 0.3009 |

**Table 5:** Statistical outcomes for models of egg/female ratios.

| Random Effects | Estimate | SE | Df | T-value | LLR | P-value |
| --- | --- | --- | --- | --- | --- | --- |
| TrtRep (Dish) | 0.00925 |  |  |  | 0.231782 | 0.6302 |
| Residual | 0.19129 |  |  |  |  |  |
| Interactions | Estimate | SE | Df | T-value | LLR | P-value |
| Temperature : pH | -0.322532 | 0.1709 | 36 | -1.887257 | 3.77374 | **0.0521** |
| Fixed Effects | Estimate | SE | Df | T-value | LLR | P-value |
| Intercept | 0.62774 | 0.08545 | 80 | 7.34626 |  |  |
| Temperature | 0.12938 | 0.12084 | 36 | 1.0706 | 0.14057 | 0.7077 |
| pH | 0.3294 | 0.12084 | 36 | 2.72582 | 3.73457 | **0.0533** |

| Random Effects | Estimate | SE | Df | T-value | LLR | P-value |
| --- | --- | --- | --- | --- | --- | --- |
| TrtRep (Dish) | 0.00455 |  |  |  | 0.564319 | 0.4525 |
| Residual | 0.05809 |  |  |  |  |  |
| Interactions | Estimate | SE | Df | T-value | LLR | P-value |
| Temperature : pH | -0.028905 | 0.098165 | 36 | -0.29445 | 0.09764 | 0.7547 |
| Fixed Effects | Estimate | SE | Df | T-value | LLR | P-value |
| Intercept | 0.30672 | 0.0489 | 79 | 6.27227 |  |  |
| Temperature | 0.19842 | 0.06967 | 36 | 2.84811 | 13.114 | **0.0003** |
| pH | 0.16569 | 0.06916 | 36 | 2.39591 | 9.36634 | **0.0022** |

**Table 6:**
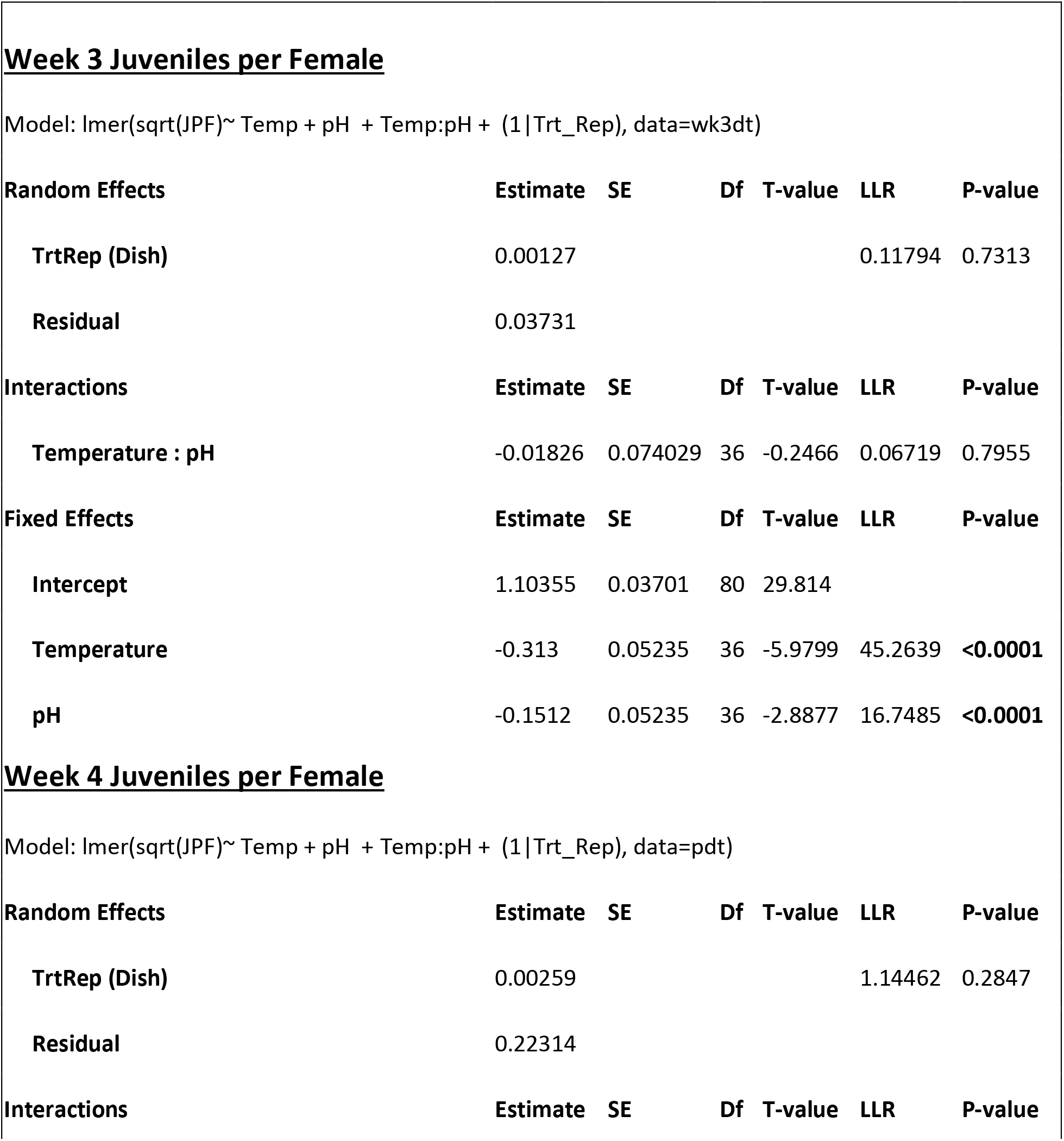
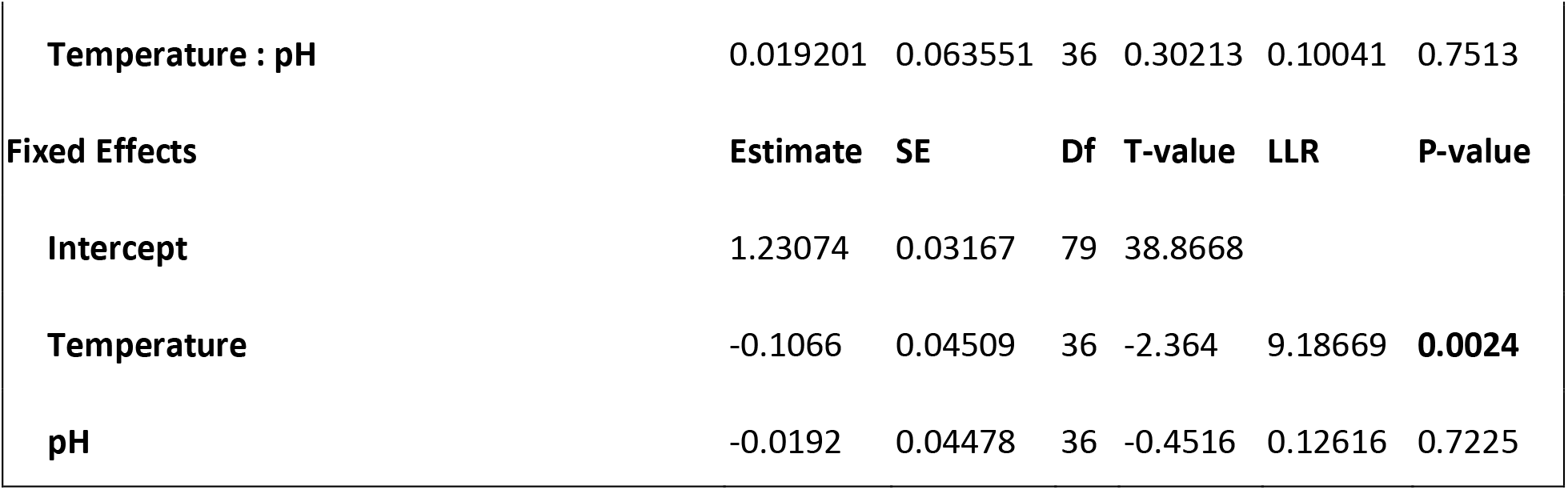
Statistical outcomes for models of juvenile/female ratios.

**Table 7:**
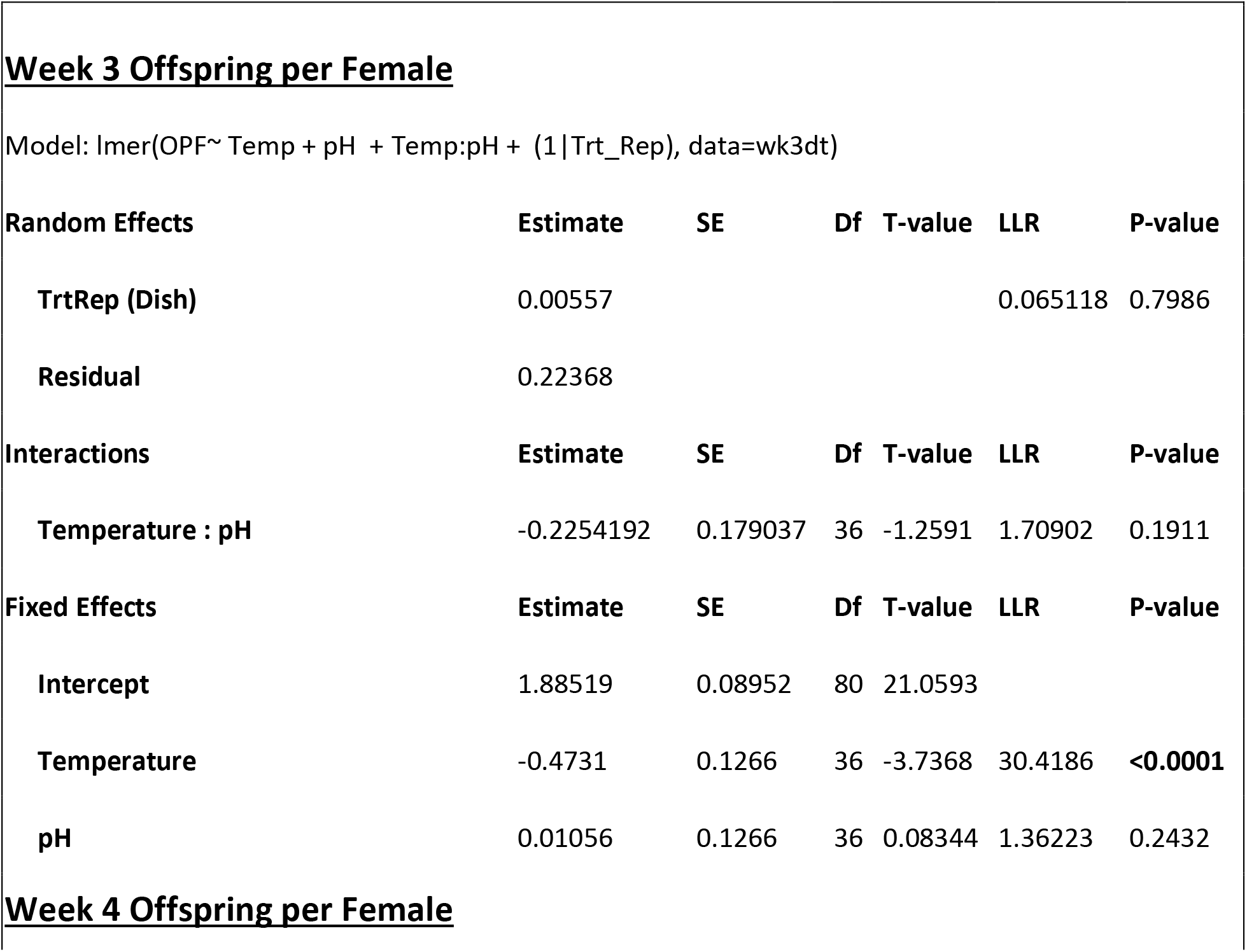
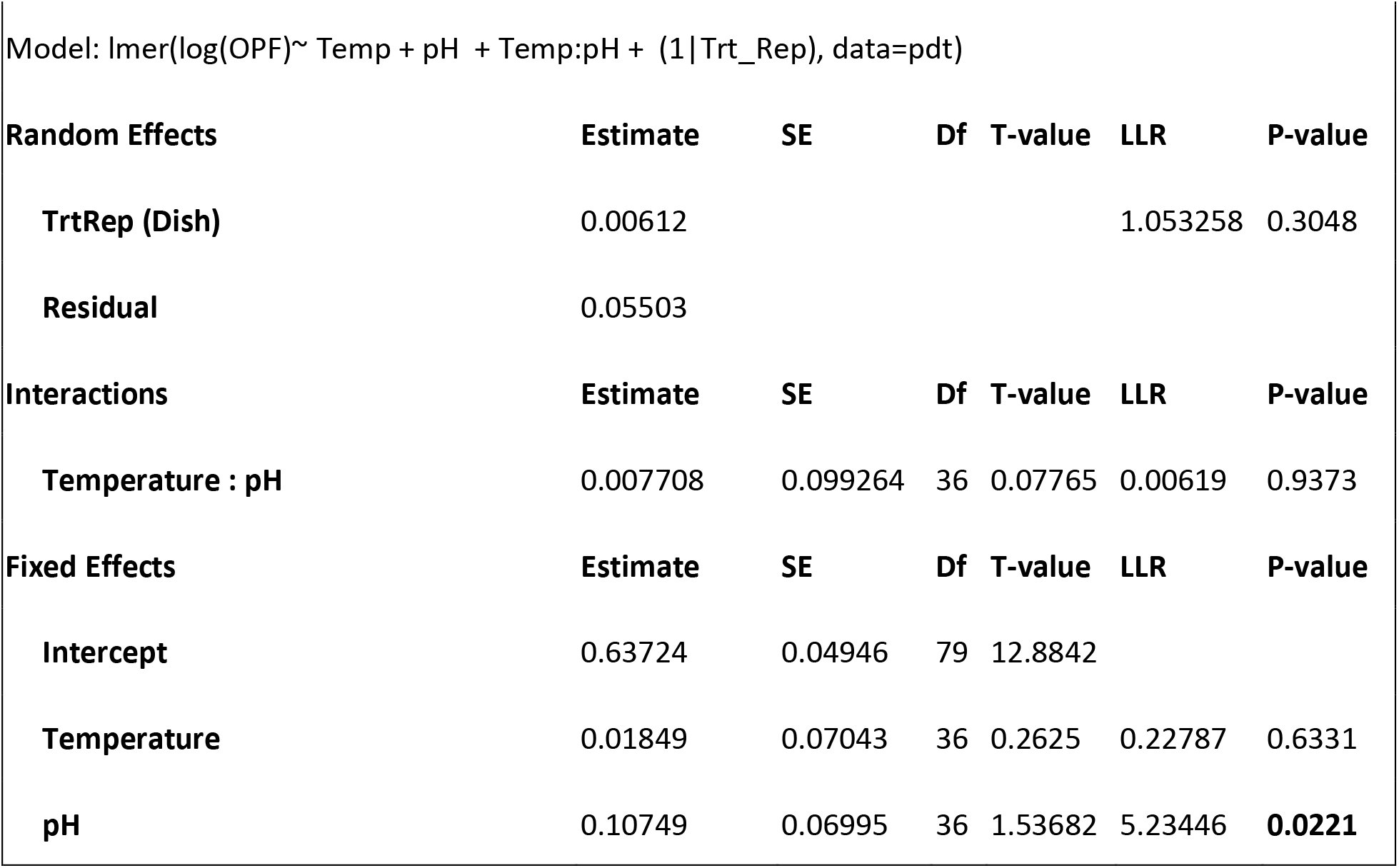
Statistical outcomes for models of offspring/female ratios.

**Table 8:** Statistical outcomes for models of ratios of proportion productive females.

|  |  |  |  |  |  |
| --- | --- | --- | --- | --- | --- |
| **Random Effects** | **Estimate** | **SE** | **z-value** | **ChiSq** | **P-value** |
| TrtRep (Dish) | 2.24E-01 |  |  |  |  |
| **Interactions** | **Estimate** | **SE** | **z-value** | **ChiSq** | **P-value** |
| Temperature : pH | -0.34443 | 0.47961 | -0.718 | 0.5117 | 0.4744 |
| **Fixed Effects** | **Estimate** | **SE** | **z-value** | **ChiSq** | **P-value** |
| Intercept | 4.05985 | 0.35058 | 11.58 |  |  |
| **Temperature** | -1.25286 | 0.37167 | -3.371 | 28.187 | **<0.0001** |
| **pH** | -0.08916 | 0.339 | -0.263 | 1.1619 | 0.2811 |

**Table 9:** Statistical outcomes for models of juvenile size data (area in um2).

| **Interactions** | **Estimate** | **SE** | **T-value** | **ChiSq** | **P-value (nested)** |
| --- | --- | --- | --- | --- | --- |
| **gammean:Temp:pH** | 0.48066 | 0.14353 | 3.349 | 6.3387 | **0.01181** |

**Subset Model for Low pH Treatments** Model: glmer(l(area\_um2/100) ~ Temp\*gammean +(1|FullTrt), family = Gamma(link="inverse"), data = wk4lph)
| **Variables** | **Estimate** | **SE** | **T-value** | **ChiSq** | **P-value** |
| --- | --- | --- | --- | --- | --- |
| **TrtRep (Dish)** | 2.93E-03 |  |  |  |  |
| **Residual** | 6.53E+01 |  |  |  |  |
| **Intercept** | 7.56E+00 | 1.86766 | 4.048 |  |  |
| temp:gammean | 0.11974 | 0.08914 | 1.343 | 1.6106 | 0.2044 |
| **temp** | -5.40444 | 20.7699 | -2.602 | 10.051 | **0.001523** |
| **gammean** | -0.03965 | 0.06543 | -0.606 | 0.3192 | 0.5721 |

**Subset Model for Ambient pH Treatments** Model: glmer(l(area\_um2/100) ~ Temp\*gammean +(1| FullTrt), family = Gamma(link="inverse"), data = wk4hph)
| Variables | Estimate | SE | T-value | ChiSq | P-value |
| --- | --- | --- | --- | --- | --- |
| TrtRep (Dish) | 1.33E-03 |  |  |  |  |
| Residual | 5.21E+01 |  |  |  |  |
| Intercept | 2.57E+00 | 1.03042 | 2.49 |  |  |
| temp:gammean | -0.36249 | 0.13335 | -2.718 | 5.3001 | **0.02132** |
| temp | 3.63827 | 1.87179 | 1.944 | 2.0815 | 0.1491 |
| gammean | 0.10426 | 0.03703 | 2.816 | 3.1812 | 0.07449 |

Subset Model for Ambient Temperature Treatments Model: glmer(l(area\_um2/100) ~ pH\*gammean +(1| FullTrt), family = Gamma(link="inverse"), data = wk4ltemp)
| Variables | Estimate | SE | T-value | ChiSq | P-value |
| --- | --- | --- | --- | --- | --- |
| TrtRep (Dish) | 4.03E-03 |  |  |  |  |
| Residual | 4.58E+01 |  |  |  |  |
| Intercept | 2.62E+00 | 1.57501 | 1.665 |  |  |
| pH:gammean | -1.46E-01 | 0.11731 | -1.246 | 2.2666 | 0.1322 |
| pH | 5.05E+00 | 3.32134 | 1.521 | 3.955 | **0.04673** |
| gammean | 1.04E-01 | 0.05653 | 1.837 | 1.1728 | 0.2788 |

| Variables | Estimate | SE | T-value | ChiSq | P-value |
| --- | --- | --- | --- | --- | --- |
| **TrtRep (Dish)** | 4.86E-04 |  |  |  |  |
| **Residual** | 8.07E+01 |  |  |  |  |
| **Intercept** | 5.95E+00 | 1.2176 | 4.887 |  |  |
| **pH:gammean** | 3.22E-01 | 0.10705 | 3.011 | 6.0391 | **0.01399** |
| **pH** | -3.90E+00 | 1.35655 | -2.874 | 0.0811 | 0.7758 |
| **gammean** | -2.40E-01 | 0.09895 | -2.427 | 0.988 | 0.3202 |

